# AXTEX-4D™: A Novel 3D *ex vivo* platform for preclinical investigations of immunotherapy agents

**DOI:** 10.1101/2020.11.05.369751

**Authors:** Ambica Baru, Swati Sharma, Biswa Pratim Das Purkayastha, Sameena Khan, Saumyabrata Mazumder, Reeshu Gupta, Prabuddha Kumar Kundu, Nupur Mehrotra Arora

## Abstract

The latest advancements in oncology are majorly focused on immuno-oncology (I-O) therapies. However, only ~7% of drugs are being approved from the preclinical discovery phase to phase 1. The most challenging issues in I-O is the problem of developing active and efficient drugs economically and on time. This mandates an urgent need for better preclinical screening models that closely mimic the *in vivo* tumor microenvironment. The established and most common methods for investigating the tumoricidal activity of I-O drugs are either two-dimensional (2D) systems or primary tumor cells in standard tissue culture vessels. However, they do not mimic the tumor microenvironment. Therefore, the more *in vivo*-like three-dimensional (3D) multicellular tumor spheroids are quickly becoming the favored model to examine immune cell-mediated responses in reaction to the administration of I-O drugs. Accordingly, we have demonstrated the utility of the three-dimensional *ex vivo* oncology model, developed on our novel AXTEX-4D™ platform to assess therapeutic efficacies of I-O drugs by investigating immune cell proliferation, migration, infiltration, cytokine profiling, and cytotoxicity of tumor tissueoids. The platform eliminates the need for additional biomolecules such as hydrogels and instead relies on the cancer cells themselves to create their own gradients and microenvironmental factors. In effect, the more comprehensive and in *vivo* like immune-oncology model developed on AXTEX-4D™ platform can be utilized for high-throughput screening of immunotherapeutic drugs.

## Introduction

Immuno-oncology therapies (I-O) have played an essential role in cancer treatment. I-O therapies reinvigorate cancer therapeutics by harnessing the immune system to direct a response towards tumors. I-O therapies cover diverse field that encompasses vaccines, checkpoint inhibitors, adoptive cell therapies, oncolytic virus therapies, and non-specific immunotherapies such as cytokines. About hundreds of clinical trials have been carried out to evaluate the efficacy of immunotherapeutic drugs[1]. However, the likelihood of success of clinical trials are very low for I-O drugs demonstrating their high failure rates[2; 3]. Moreover, effectiveness of I-O therapies is not same for all cancer patients and types. Developing active drugs economically and in a timely manner is one of the biggest challenges in oncology. Considering the failure rate, high cost and time-consuming nature of the clinical development of oncology drugs, there is an urgent need for improved oncological models for drug screening that more closely resemble and mimic the tumor microenvironment *in vivo*. The crosstalk between tumor cells and other immune cells present in the tumor microenvironment, such as T cells, plays an important role in tumor progression. 3D models have been shown as valuable models to understand the underlying mechanisms of immune cell interaction with the tumor microenvironment [4] These models establish novel therapeutic concepts in the modern area of immune-oncology and could prevent or at least reduce therapy failure in the future by the screening of immunotherapeutic drugs.

Traditionally, tumoricidal activity and immune evasion have been evaluated in two-dimensional (2D)-systems. However, 2D systems do not simulate the 3D microenvironment of the original tumors, and therefore, anticancer drugs tested on these systems are eliminated during clinical development. The tissue-specific architecture involving elements of the surrounding microenvironment such as stromal cells are essential components of a tumor and can be at least partially recapitulated utilizing three-dimensional (3D) cell culture models. The stromal cells not only provide a scaffold for tumor growth, but they also participate actively in tumor formation, progression, and metastasis; produce cytokines/chemokines, and help in the manufacture of the extracellular matrix. Thus 3D-culture systems better simulating the *in vivo* tumor microenvironment have received attention to avoid certain drawbacks of 2D-culture models [5; 6; 7; 8].

The procedures in place for the development of drugs involve thorough evaluation of novel drug candidates in both pre-clinical and clinical phases by evaluating their immune mediated responses. In *in vivo* conditions, the immune cells need to infiltrate the 3D cell-matrix formation to attack the malignant target cells[9; 10]. Therefore, they need to overcome physical barriers. The structural complexity of spheroids by providing physiological barriers to immune cells allows for higher resistance to cytotoxicity. Thus, 3D cell culture provides more physiological disease modelling and may ultimately result in improved translation and a reduction in number of the animal models required in drug discovery or screening programmes.

Improving 3D oncology models will create screening tools with greater accuracy in assessing therapeutic efficacies of I-O drugs. To this end, the 3D tumor tissueoids were generated in less than 72 hrs on our novel AXTEX-4D™ platform[11], which helps create an extracellular and intracellular architecture of tissueoids and thus emulate tumor microenvironment(TME). This study demonstrates the utility of a novel 96 well 3D *ex vivo* tissueoids model to screen I-O drugs in a cost and time-effective manner by studying cancer/immune cell interactions to investigate immune cell-mediated responses such as migration, infiltration, tumor cytotoxicity, and tumor immune evasion.

## Results

### T-cells mediated immune responses and 3D cytotoxicity

The success of T cell-based immunotherapy is closely associated with the activation, ability of effector T cells to migrate to the target area, and making contact with their malignant targets[12]. To address this, we first confirmed the presence and activation of blood T-cells by providing co-stimulatory signals with anti CD3 and anti CD28 antibody for 2,4,5 and 7 days with 1-day culture taken as reference and counted 100 percent. Unstimulated T-cells were taken as a negative control. Flow cytometry confirms the presence of T-cells, which were isolated from human PBMCs (Figure 1A; T-cells=72.66±0.62%; n=3). Compared to unstimulated T-cells, differential morphology was observed in stimulated T cells (Figure 1B). The proliferation of T-cells was highest on day 4 (2.2±0.06 fold vs. day 1; p<0.05) as compared to 2(1.8±0.006 fold vs. day 1), 5(2.1±0.18 vs. day 1), and 7 (1.2±0.30 fold vs. day 1) days (Figure 1C). However, unstimulated T cells did not show any significant change in proliferation over time, suggesting that T cells’ stimulation is essential for an immune response (2 days:0.69±0.07 fold, 4 days: 0.78±0.09 fold, 5 days:0.54±0.09 fold, 7 days: 0.39±0.05 fold). Therefore, all the future experiments were done with 4th day culture of stimulated blood T cells. An important component of any immune response is the migration, cytokine production, and infiltration of immune cells to the target area requiring defense. These processes play a significant role in accessing the efficacy and success rates of I-O drugs. To evaluate the migration, IFN-γ secretion, and cytotoxic effects of immune cells on 3D tissueoids, T cells were co-cultured with MCF-7 tissueoids. The number of migrated T cells towards MCF-7 tissueoids and IFN-γ secretion was highest at 48hrs (Migration: 50.25±0.22%;p<0.05; IFN-γ:0.91±0.002 ng/ml) in comparison to 24 (Migration: 16.5±0.68%; IFN-γ: 0.59±0.02 ng/ml) and 72hrs (Migration:6.75±0.64%; IFN-γ:0.05±0.0008 ng/ml) (Figure 1D-1E). No significant difference in migration and IFN-γ secretion was observed with unstimulated T-cells. As expected, compared to control (only MCF-7 tissueoids), the proliferation was maximally inhibited in co-cultured tissueoids at 48hrs (T cells+MCF-7 tissueoids-48hrs:1.27±0.02 fold; 24hrs:1.85±0.012; 72hrs: 1.55±0.11 fold), supporting the specific targeting of the T cells to the MCF-7 tissueoids (Figure 2A). Since the maximum migration, IFN-γ production, and cytotoxic response was observed at 48hrs, we next access the infiltration of unstimulated/stimulated T cells with MCF-tissueoids at this period. Stimulated T-cells showed significantly higher infiltration in comparison to unstimulated T cells at 48hrs (Figure 2B). These studies suggest that the immune T cells cannot just reach, infiltrate, and kill the target tumor tissueoids grown on AXTEX-4D™ platform but can also produce IFN-γ, a cytokine primarily involved in host defence, immune surveillance, and the establishment of adaptive immunity[13].

**Figure 1:**
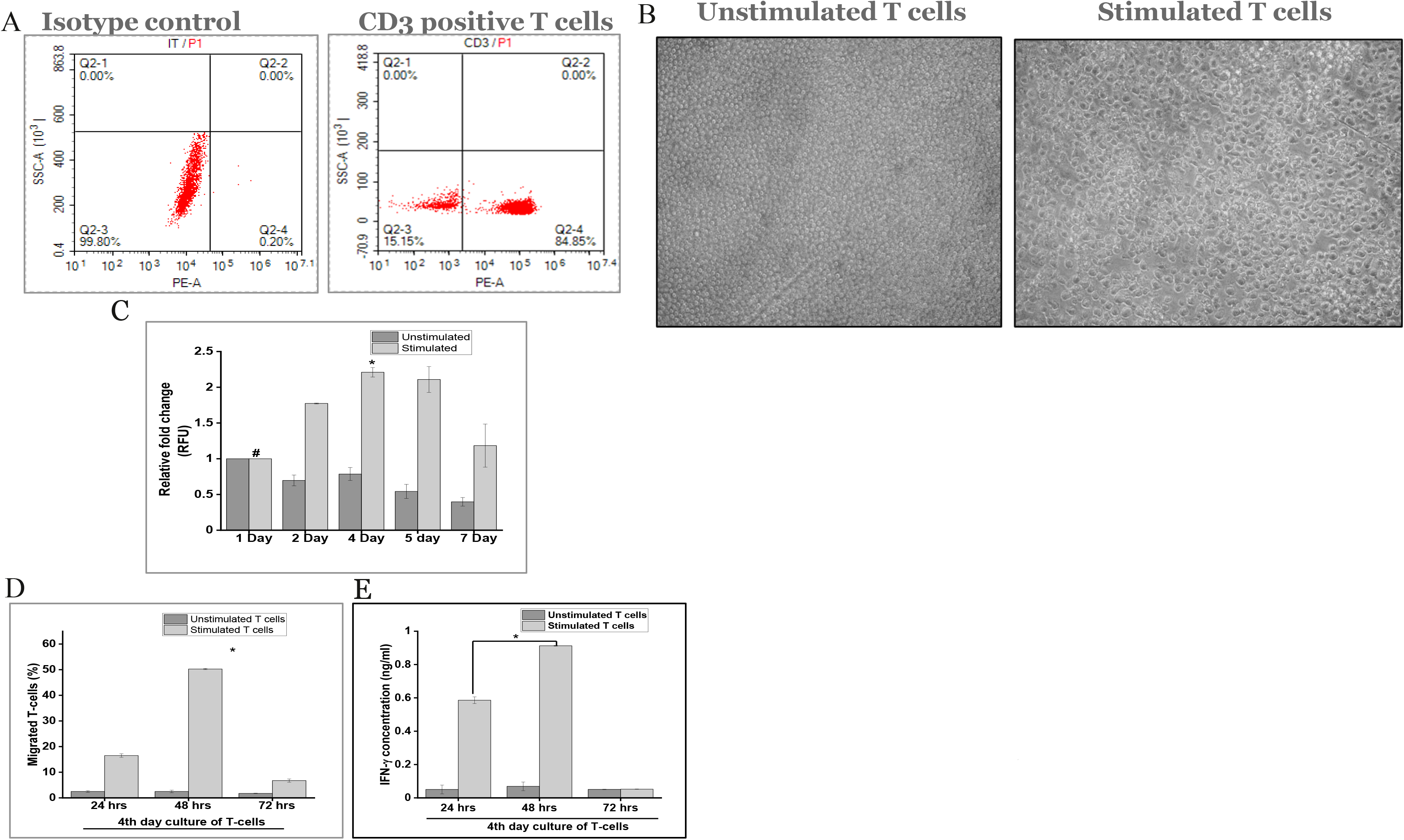
Elucidation of T-cell mediated Immune response against breast cancer tissueoids (MCF-7) grown on AXTEX-4D™ platform. A) Confirmation of T-cells in human PBMCs by flow cytometry B) T-cells were stimulated with anti CD3 and anti CD28 antibody. Bright field microscopy images for unstimulated (left panel) and stimulated T cells (right panel) C) Prestoblue assay depicting proliferation of stimulated and unstimulated T cells at the indicated time periods. D) Migration and E) IFN-γ production by 4th days culture of unstimulated and stimulated blood T cells when co-cultured with MCF-7 tissueoids. Results from one representative donor is shown here. Graph bars summarized the results of three independent experiments. Values are means ± S.E.M

**Figure 2:**
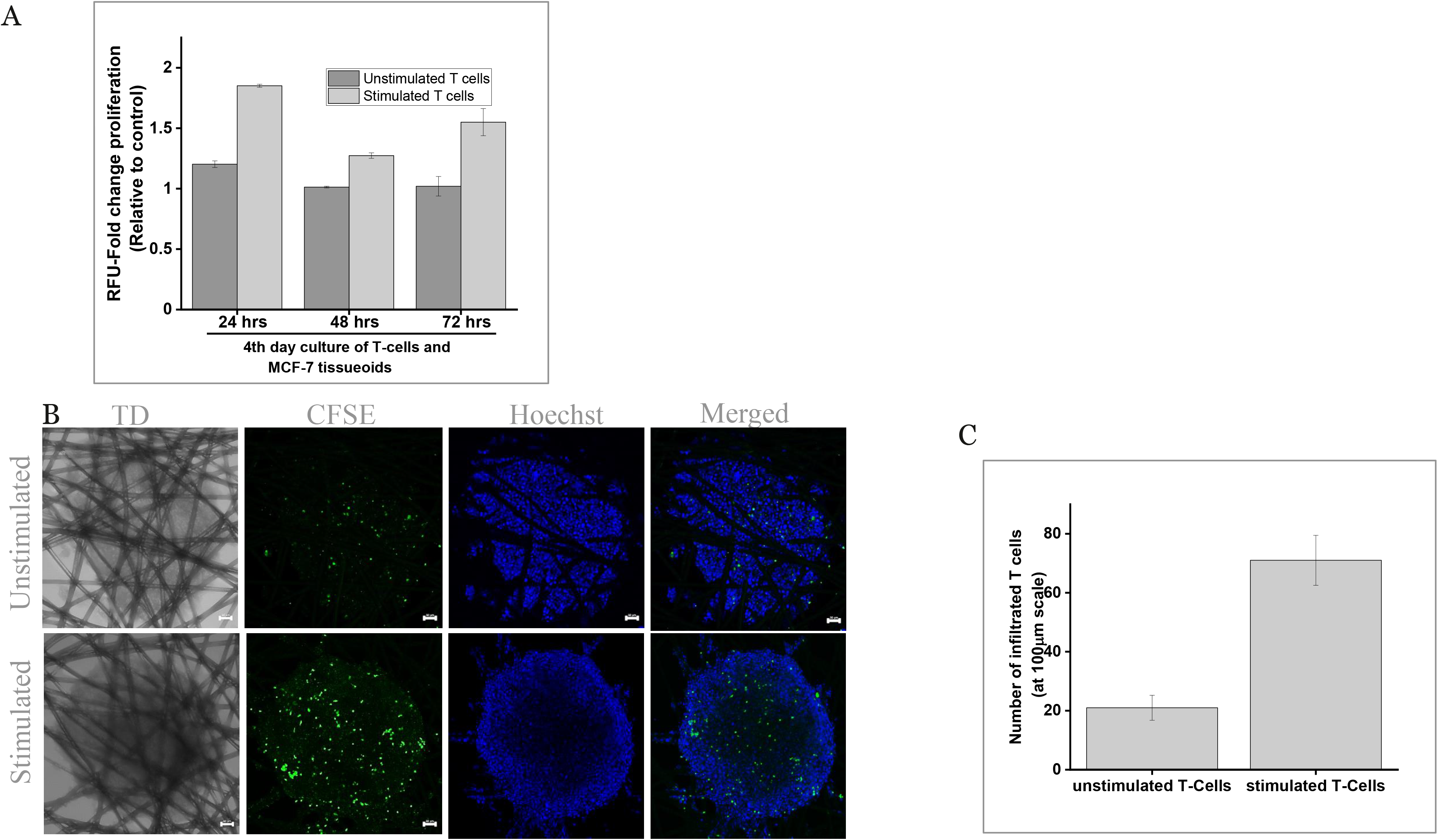
T-cell mediated cytotoxicity and infiltration of blood T cells towards MCF-7 tissueoids. A) Proliferation data given as the fold change in RFU, measured using PrestoBlue at the indicated time periods B) MCF-7 tissueoids (3D) were co-cultured with unstimulated and stimulated T-cells for 48hrs and stained with Hoechst (blue). Blood T (green) were stained with CFSE green. Confocal microscopy showed noticeable interspersion of stimulated blood T-cells (lower panel) with the blue MCF-7/Hoechst cells of the tissueoids as compared to unstimulated T cells (upper panel) C) Bar chart representation of infiltrated T-cells. Error bars indicate SEM)

### Effect of chemo attractants on the infiltration of Jurkat T-cells

Immune cell recruitment to the target area and their infiltration has been shown to correlate in increasing immunotherapeutic drugs’ efficacy and thus increased patients’ survival. 3D models have been shown as valuable models that are capable of establishing novel therapeutic concepts in the area of immune-oncology[14]. To observe the effect of chemo attractant on immune cell infiltration into the MCF-7 tissueoids, Jurkat cells were allowed to migrate overnight toward medium containing SDF-1α or 10% FBS, a known chemo attractants for lymphocytes[15]. Confocal microscopy showed noticeable interspersion of CFSE labelled Jurkat-cells with the MCF-7 tissueoids (Figure 3A). Infiltration of Jurkat cells was found to be significantly higher in the presence of 10% FBS as compared to infiltration observed with SDF-1α and 0% FBS (scale: 10μM-10%FBS: infiltrated T-cells-99±1.41; SDF-1α: 28±8.48; 0%FBS: 9±4.24), (Figure 3-B). These studies suggest that the AXTEX-4D™ platform could be used for evaluating conditions which enhance or suppress immune cell infiltration to target area.

**Figure 3:**
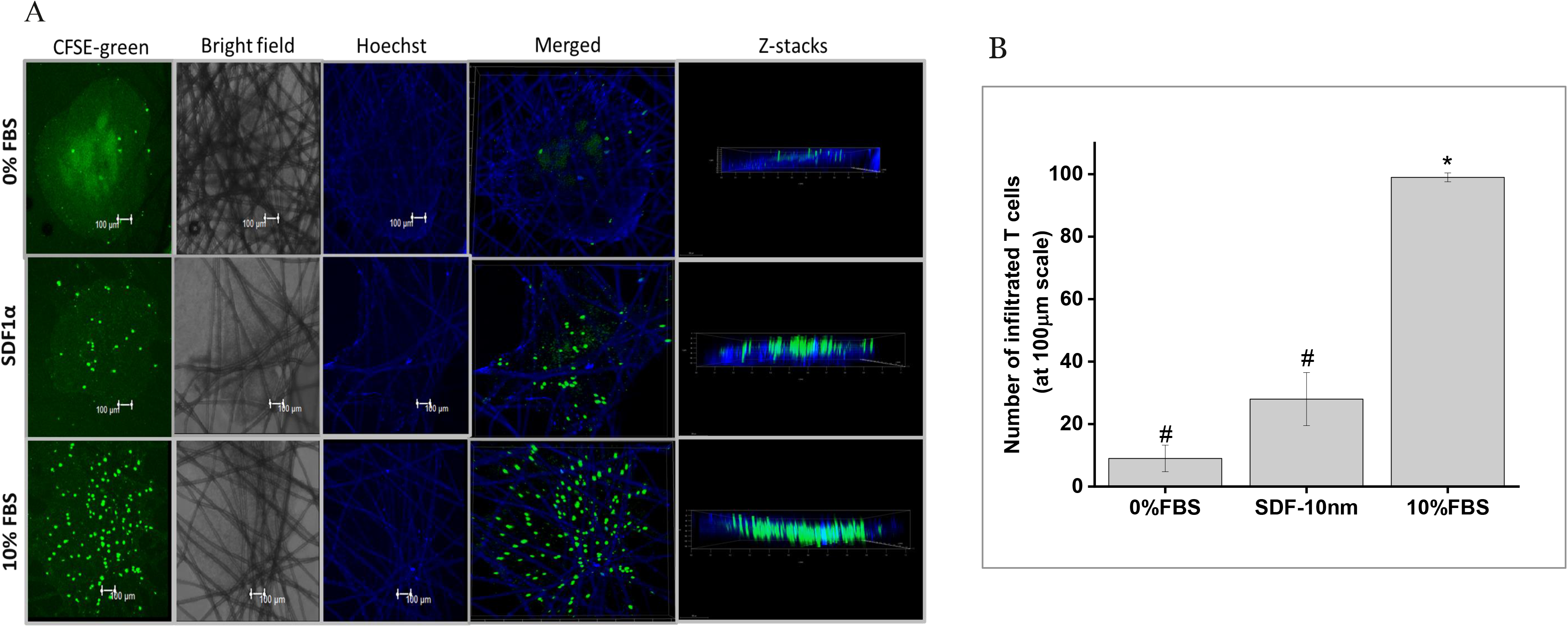
Infiltration of Jurkat cells towards MCF-7 tissueoids in the presence of chemo attractants. Confocal microscopy images depicting infiltration of Jurkat cells withMCF-7 tissueoids in the presence of SDF-1α and 10%FBS. B) Bar chart representation of infiltrated T-cells. Error bars indicate SEM.

### Immune response of 3D tissueoids against immune checkpoint inhibitors

Knowledge about PD-L1 expression needs to be put into the context of the PD-1 expression by T cells[16] to observe immune-suppressive activity of tumor cells[17; 18]. Therefore, we first aimed to measure the percentage of PD-L1 and PD-1 expressing cells in tumor tissueoids (MCF-7 and A-375) and T cells, respectively. (Figure 4A-B). The surface expression of PD-L1 was determined by flow cytometric analysis. About 18.5±2.12% of A-375 cells were positive for PD-L1. However, less than 1% of MCF-7 cells (0.41±0.21%) were found to be positive for PD-L1, suggesting negligible expression of PD-L1 by these cells when compared with A-375 cells (P<0.001; Fig. 4A-B). Similarly, 22±0.53% of activated T-cells were positive for PD-1. The PD-1/PD-L1 blockade therapy induces apoptosis of tumors by enhancing T lymphocyte proliferation, cytokine production, and survival. To determine whether PD-L1/PD-1 blockade therapy can induce the immune response in 3D tissueoids, we first blocked the PD-L1/PD-1 interaction in two ways: 1) treating PD-L1 expressing A-375 tissueoids with 2.5 μg/ml of Atezolizumab (AL-2.5) 2) treating blood T-cells with 25 μg/ml of Nivolumab (NP-25). Flow cytometry confirmed that detection of the PD-L1 and PD-1 epitope was blocked by the specific antibodies after 30minutes of treatment in A-375 and T cells, respectively (AL-2.5: 1.22±0.55%; NP-25: 0.32±0.21%) (Figure 4B-D). Maximum IFN-g production was observed in AL-2.5 treated A-375 tissueoids (2.87±0.013) when compared with untreated T-cells/A-375 tissueoids (1.27±0.007ng/ml) and NP-25 (NP-25: 1.47±0.005ng/ml) (Figure 5A). In comparison to control, an obvious increased cell proliferation of T-cells/A-375 tissueoids (T-cells/A-375 tissueoids:1.46±0.02 fold) speculate their immune-resistance nature. AL-2.5 treatment significantly decreased cell proliferation in T-cells/A-375 tissueoids compared to untreated co-cultured tissueoids (AL-2.5:0.88±0.006 fold; p<0.05). However, the difference was not significant with NP-25 treatment (NP-25: 1.18±0.02 fold)(Figure 5C). As expected, the treatment of AL-2.5 and NP-25 did not lead to any significant effect on IFN-g production or cytotoxicity of MCF-7 tissueoids (Figure 5B, D). These findings suggest that oncology model developed on AXTEX-4D™ platform cannot just distinguish T-cells response in tumor tissueoids having differential expression of immune checkpoint biomarkers (PD-L1) but could also be used to evaluate the efficacy of immunotherapeutic drugs.

**Figure 4:**
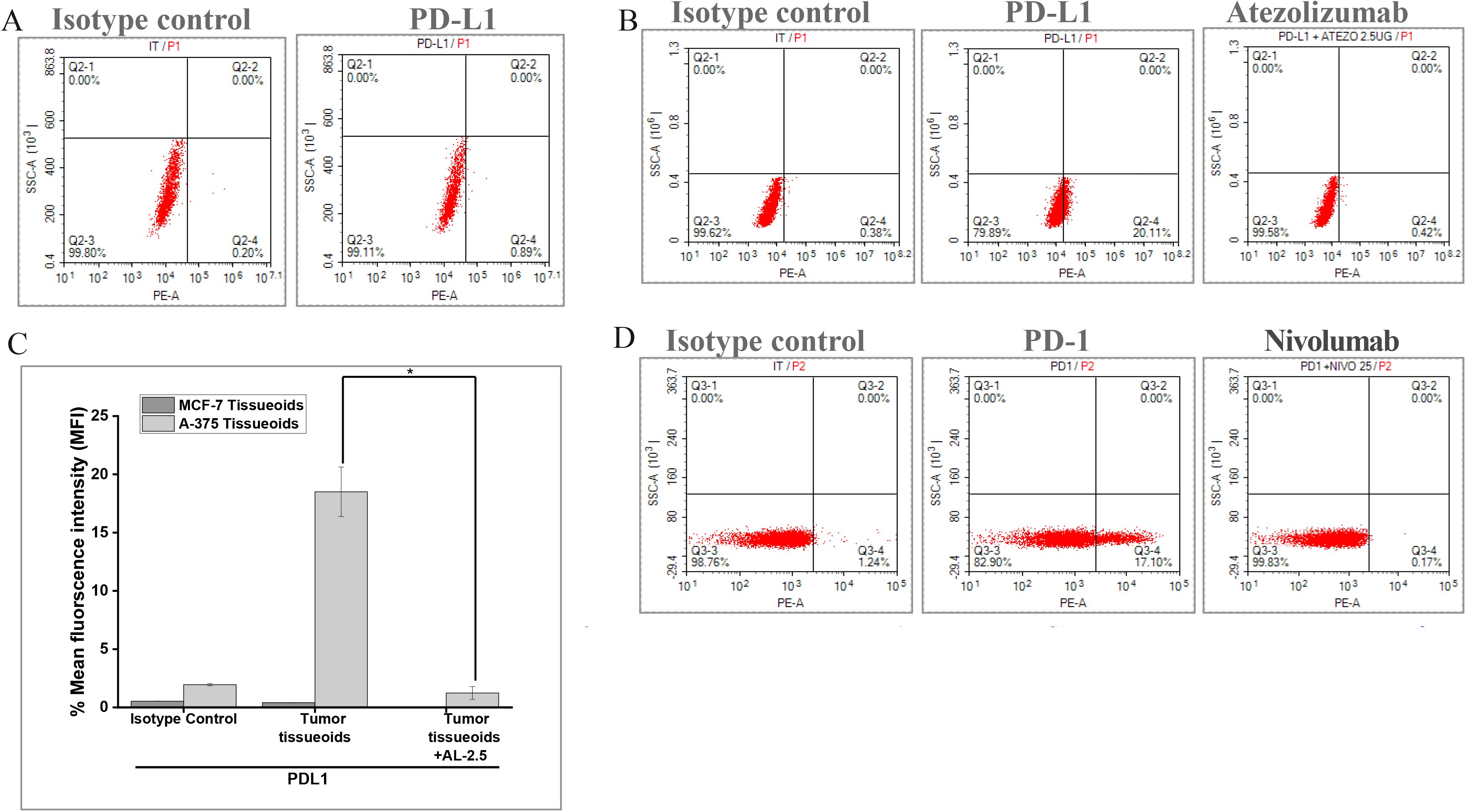
Blocking activity of Atezolizumab and Nivolumab in tissueoids grown on AXTEX-4D™ platform. Percentage of PD-L1 expressing cells in A) MCF-7 and B) untreated (middle panel) and atezolizumab (AL-2.5) treated A-375 tissueoids (right panel). The surface expression of PD-L1 was determined at 30 minutes post treatment by flow cytometry. C) Bar chart representing % MFI in AL-2.5 treated or untreated tissueoids D) Percentage of PD-1 expressing cells in untreated(middle panel) andNivolumab treated blood Tcells (right panel)

**Figure 5:**
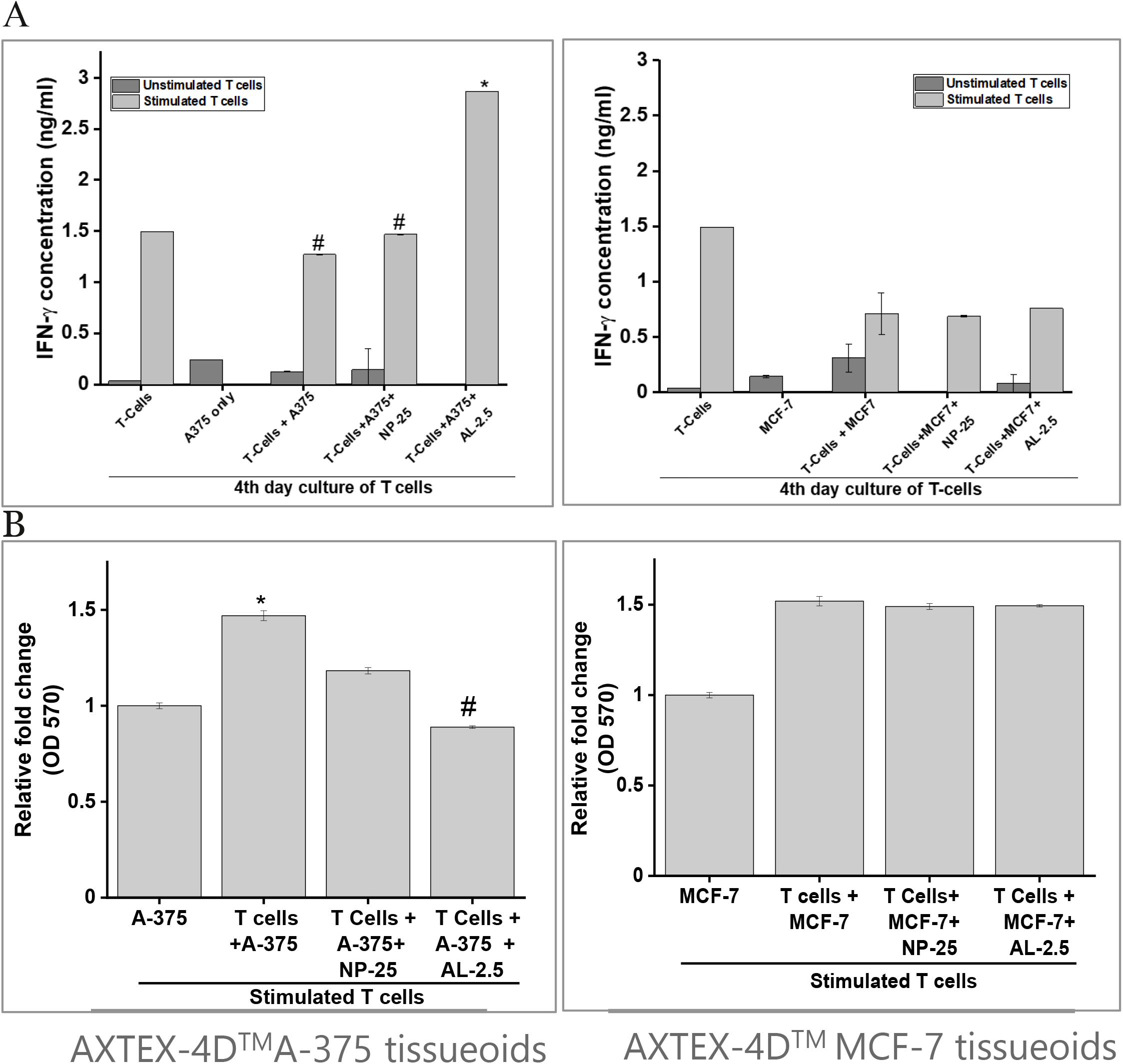
Immune response of tissueoids against immune checkpoint blockade therapy. A) Tissueoids were treated with 2.5μg/ml of Atezolizumab (AL-2.5) and T-cells were treated with 25μg/ml of Nivolumab (NP-25). ELISA depicting IFN-γ production by the treated and untreated T-cells when co-cultured with A-375 (left panel) and MCF-7 tissueoids (right panel). B) Effect of NP-25 and AL-2.5 on proliferation of two tissueoids were studied by MTT assay. Values are means ± S.E.M.

### Drug sensitivity in 2D- and 3D-culture

PD-L1 blockade therapy by blocking interaction of PD-1 and PD-L1 leads to activation-induced cell death of tumors. However, overall, only a small percentage of patients currently benefit from PD-L1 blockade therapy. Many mechanisms in tumor cells help in acquiring resistance to immune checkpoint blockade (ICB) therapy[19]. Previous studies have shown that cells grown in 3D models showed an increased resistance to I-O drugs as compared to 2D model[20; 21]. To evaluate this, we first compare the immune resistance feature of cancer cells in 2D vs. 3D tissueoids model. In comparison to controls, we observe a significant increase in cell viability (1.46±0.02 fold vs 3D A-375 controls) and comparatively low IFN-g secretion of 3D A-375 tissueoids when co-cultured with activated T-cells (A-375-cells: 1.5±0.0001ng/ml; T-cells/A-375 tissueoids:1.2±0.006ng/ml). These results suggest that the developed model is capable of regulating mechanisms associated with immunosurveillance against tumors. However, an obvious decrease in cell viability (0.08±0.013 fold vs. 2D A-375) and relatively higher IFN-g secretion (A-375 cells: 1.41±0ng/ml; T-cells/2D-A375:1.7±0.003ng/ml) of 2D A-375 cells speculates lacking specific immune resistance features of *in vivo* primary tumors. We speculate that the overestimated cytotoxicity, as observed in 2D culture system, may not be able to differentiate appropriate response against I-O drugs[22; 23]. To evaluate this, next, we observed the effect of AL-2.5 and NP-25 on cytotoxicity and cytokine production by A-375 tissueoids and 2D cultures of A-375 cells. Treatment with AL-2.5 comparatively reduce cell proliferation (AL-2.5 treated T-cells/A-375: 0.88±0.006-fold; untreated T-cells/A-375: 1.46±0.02 fold) in T-cells/ A-375 tissueoids, while no significant difference was observed with the 2D model (Figure 6A). This suggests that blockage of PD-1-PD-L1 interaction T cells can induce cell cytotoxicity in A-375 tissueoids. AL-2.5 treatment enhance the production of IFN-g in co-cultured cells (T-cells/A-375 tissueoids+AL1-2.5: 2.4±0.04 ng/ml) as compared to the untreated control cells (T-cells/A-375 tissueoids: 1.2±0.006 ng/ml; p<0.05) (Figure 6B). These findings support the previous studies demonstrating the role of PD-1-PD-L1 checkpoint blockade therapy in the upregulation of IFN-γ, which in turn plays critical roles in the clearance of tumor cells[24; 25]. However, no such difference was observed with 2D model (T-cells/2D A-375 +AL-2.5:1.57±0.06;T-cells/2D A-375: 1.63±0.03 ng/ml; p<0.05). These data speculate the role of 3D-culture in determining the sensitivity of the cell lines to I-O drugs by mimicking the tumor microenvironment, while no such difference was observed in the case of 2D culture model.

**Figure 6:**
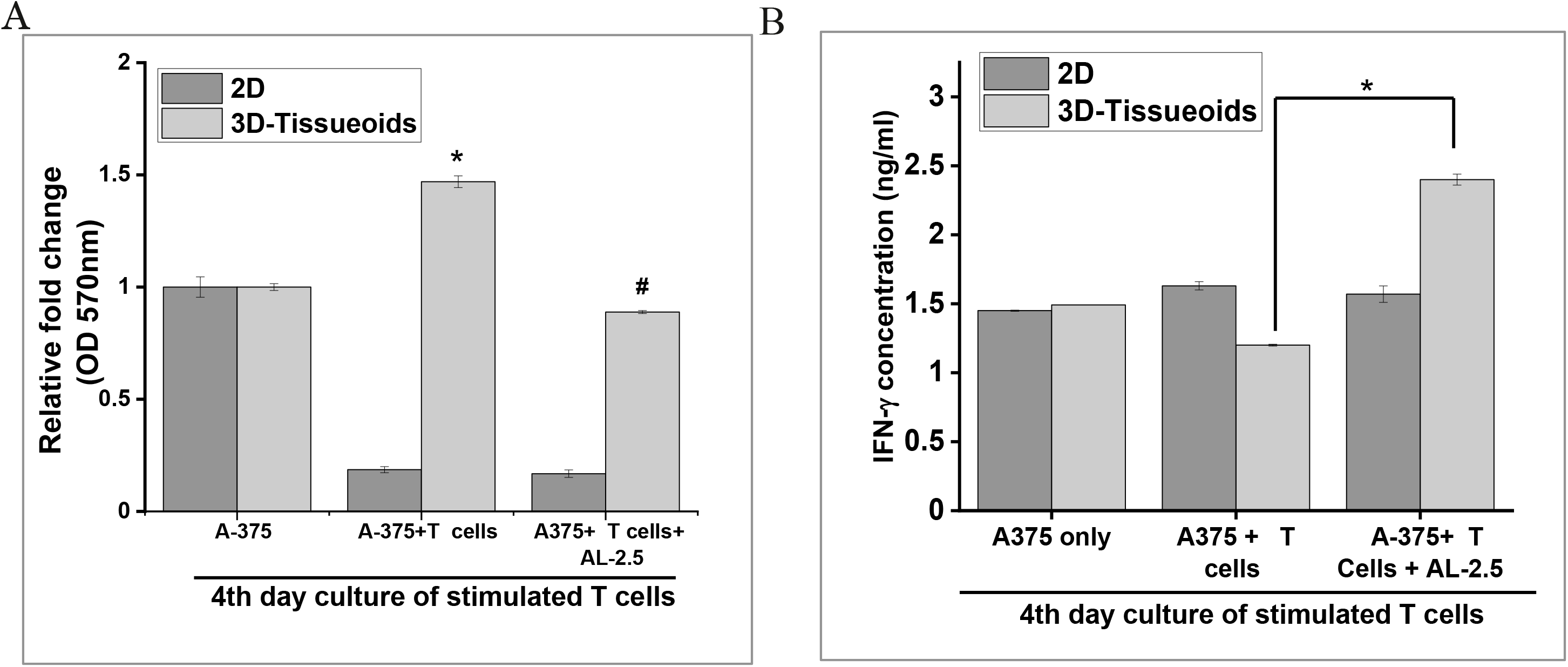
Effect of Atezolizumab (AL-2.5) on IFN-γ secretion and cell viability in 2D and 3D tissueoids model. A-375 cells were grown on either 2D or 3D tissueoids followed by treatment with AL2.5 (2.5μg/ml) for 48 hrs. A) Cytotoxicity was observed by MTT assay. B) ELISAdepicting IFN-γ production by the treated and untreated T-cells when co-cultured withA-375 tissueoids. Values are means ± S.E.M.

**Figure 7:**
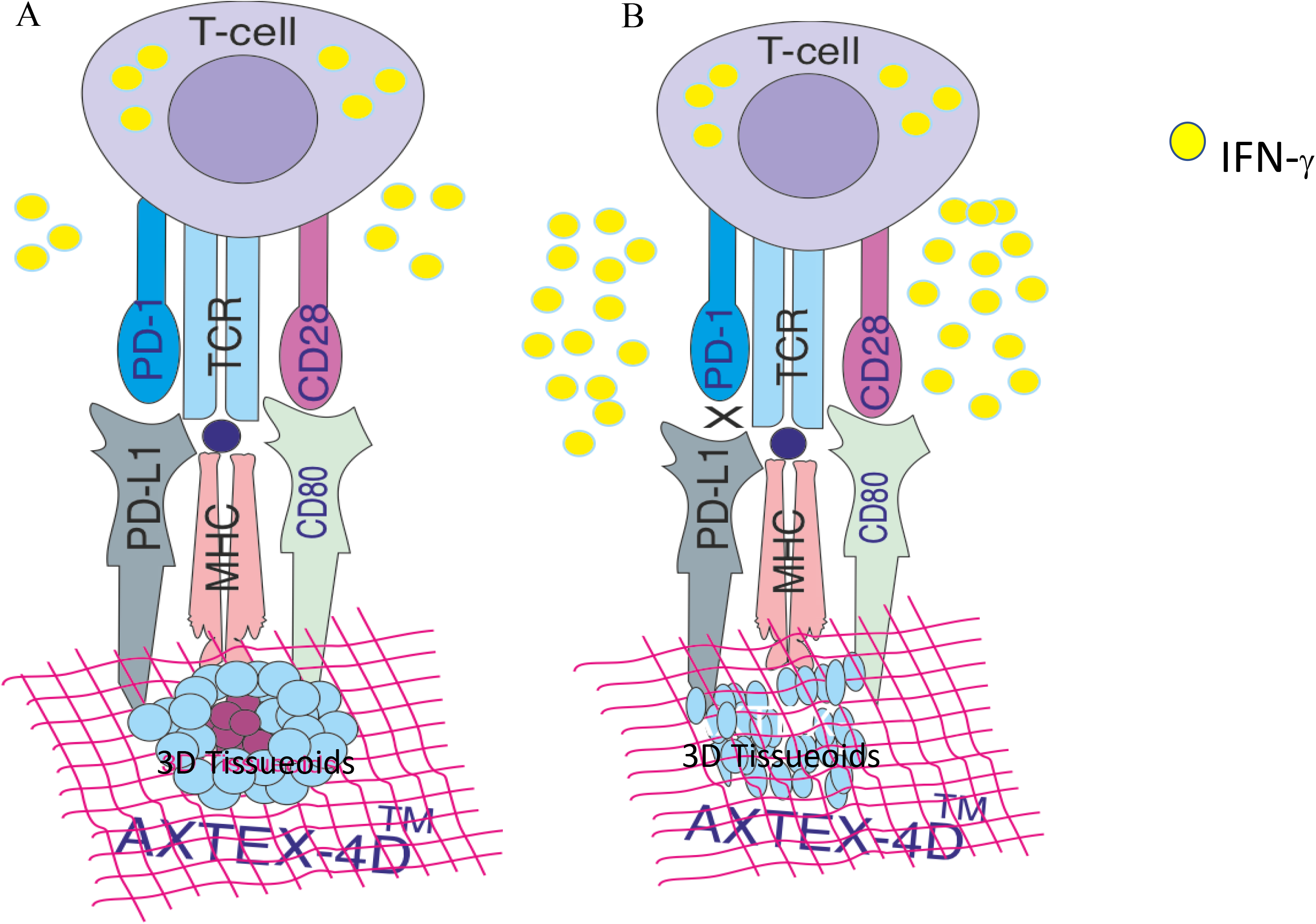
Schematic representation of Immune checkpoint blockade therapy in cancer tissueoids: A) PD-L1 signals through T-cell PD-1 to “turn off” T cells and IFN-γ secretion in order to minimize damage to cancer tissueoids. B)Blocking the PD-1–PD-L1 pathway by using either a PD-1 antibody or a PD-L1 antibody allows for T-lymphocytes to mount a robust immune response against tumor tissueoids.

## Discussion

Currently, as many as hundreds of clinical trials are being carried out globally to evaluate the efficacy of multiple immunotherapeutic drugs as monotherapy or in combination[1].However, the failure rate of these drugs is quite high due to the lack of clinical efficacy and/or unacceptable toxicity[2; 3]. Concerning the high attrition rates of clinical trials[2; 3], there is an urgent need to develop a suitable tumor model for faster and reliable screening of immunotherapeutic drugs. 3D models play an important role in identifying therapeutic candidates since they at least partially mimic the tumor microenvironment. In scaffold -based 3D cultures, scaffolds can be biological (hydrogels such as collagen, gelatin, alginate, or chitosan) or synthetically engineered (Polyethylene glycol (PEG), polylactic acid (PA), polyglycolic acid (PGA)) to emulate key properties of ECM. Whereas, natural hydrogels are weak by nature, synthetic hydrogels are often of poor biocompatibility and produce non-natural degradation products[22]. These products could be toxic and pose a potential risk if used in biomedical applications. Additionally, these systems are not entirely efficient in being tedious to produce, time-consuming, unstable over a long period, and may pose sample retrieval challenges[23]. The double network hydrogels have attracted much attention. Still, they often contain a chemical crosslink formed by UV curing or a chemical process that might be unfavourable to cells[24]. Hence, developing a robust hydrogel for both bioprinting and cell culturing is still a challenge[25]. This study demonstrates the significance of a ex vivo AXTEX-4D™ platform in investigating immune cell migration, infiltration, cytokine release, and 3D tumor cytotoxicity in a single, high-throughput 96 well format. The platform is non-influencing and relies on the tumor cells to create their microenvironment naturally.

An important component of any immune response is the activation, migration, and subsequent infiltration of immune T cells to the target area requiring defence. Tumor and its microenvironment influences T cell trafficking and function and posed obstacles in T-cell therapy[26]. Therefore, in the absence of a model that closely mimics the tumor microenvironment, it is hard to imagine the success rate of immunotherapeutic drugs. The drugs may bind to their target but may not activate an immune response. This study showed that blood T cells were able to migrate, infiltrate and induce cytotoxicity in 3D MCF-7 tumor tissueoids. We speculate that AXTEX-4D™ tumor model provided an added structural complexity level to investigate 3D tumor cytotoxicity and tumor immune evasion. These results were closely matched with the previous findings where 3D spheroids were shown to closely mimic the tumor microenvironment and recapitulate tumoricidal activity seen *in vivo*[10; 27; 28; 29].

Cancer cells are capable of activating different immune checkpoint pathways that harbour immunosuppressive functions. However, only a few patients respond to these immune checkpoint therapies due to tumor cells’ immune resistance. Tumor spheroids were found to express PD-L1 ligands, making them a target for T-cell mediated destruction due to the PD1-PDL1 interaction. Tumor spheroids can escape the immune mechanism by shedding PDL1 receptor ligands[30]. Reports suggest that knowledge about PD-L1 expression needs to be put into the context of the presence or absence of a T cell infiltrate by the blockage of PD-1 engagement[16]. This knowledge would help further interpret the meaning of the assay results and provide insights into the mechanisms of cancer escape from immune surveillance and the potential for response to ICB therapy. PD-1/PD-L1 blockage by inhibitors was capable of increasing IFN-γ secretion, cytotoxicity for A-375 tissueoids (PD-L1 positive), while, as expected, the difference was not found to be significant in MCF-7 tissueoids (PD-L1 negative) when compared with untreated co-cultured cells. These findings suggest the significance of AXTEX-4D™ platform in evaluating the efficacy of immunotherapeutic drugs on the basis of differential expression of biomarkers by tissueoids.

Previous studies speculate that the development of anticancer drugs based on screening of T-cells migration using 2D-cultured cell lines is not efficient[31; 32]. Standard human cell culture models do not recreate the 3D, multicellular interactions that direct the immune response against cancer or the tumor cell-immune cell interactions that dictate immunotherapies’ success[33]. Our present study demonstrates that tissueoids grown on AXTEX-4D™ platform are more resistant to cytotoxicity caused by activated T-cells in a 3D-than in 2D-culture. In contrast, the efficacy of immunotherapeutic drugs could be validated more precisely in 3D culture than 2D culture. These findings support the previous findings where numerous anticancer drugs are eliminated during clinical development due to the overestimated anticancer activity of therapeutic drugs on a 2D-culture-based screening platform[10; 15].

This study aims to demonstrate that the *ex vivo* tumor tissueoids model developed on AXTEX-4D™ platform could be amenable to assay numerous different targets, effector cell types, immune responses and is a powerful tool for evaluating and screening immunotherapeutic drugs.

## Materials and Methods

### Cell Lines and Primary Tissues

The two human cancer cell lines (MCF-7 (Breast Cancer) and A-375 (Skin carcinoma cells) were obtained from ATCC. MCF-7 was cultured in EMEM (Sigma-Aldrich, St; Louis, MO, USA), while A-375 cells were cultured in DMEM (Sigma-Aldrich, St; Louis, MO, USA). All the adherent cell lines were cultured in presence of 10% FBS (Gibco) and supplemented with 2 mM glutamine (Sigma-Aldrich, St; Louis, MO, USA). Cells were cultured at 37°C humidified condition with 8% CO2 under static condition.

### Donor information

Blood from the healthy donors was collected for isolation of T-cells from PBMCs. The study is approved by the Institutional Ethics committee at Premas Biotech Pvt. Ltd.

### T-cell Isolation

The Pan T Cell Isolation Kit II (Miltenyi Biotec, USA) was used for indirect isolation of T cells (CD3+CD8+ cytotoxic T cells and CD3+CD4+ T helper cells) from human peripheral blood mononuclear cells (PBMCs). Briefly, 10 ml EDTA blood was collected from donor and layered on top of the histopaque. The blood was centrifuged at 400 g for 20 mins at 22°C. After centrifugation, cells were isolated from buffy coat appeared between plasma and histopaque and washed twice with 1X PBS. Next, cells were incubated for 5 minutes with 10 μl of PAN T cells Biotin antibody cocktail and 20 μl of PAN-T cell microbead cocktail for 10 min at 2-8°C. Finally, cocktail was passed through the column. The unlabelled T cells were collected in the pass through and centrifuged at 600 rpm for 5 min. The pellet was resuspended in flow cytometry staining buffer (FACS) buffer containing 1:100 dilution of CD3 antibody (Cat≠317307; Biolegend) and incubated for 30 minutes at 4°C in the dark. After centrifugation and two times washing with FACS buffer, data were acquired using ACEA Novo flow cytometer

### Activation of T cells

96-well plate was coated with 5 μg/mL solution of anti-CD3 antibody in sterile PBS. Just before adding cells, removed the antibody solution and rinsed plate with PBS two times to remove all unbound antibody from each well. Following which, 100 μL of the cell suspension was added to each well with soluble anti-CD28 antibody at 2 ug/ml and incubated the cells in a humidified 37°C, 5% CO2 incubator for 2,4,5 and 7 days. For the control unstimulated cells, wells were coated with sterile PBS and kept at 4°C overnight.

### AXTEX-4D™ Scaffold

It is a non-woven fabric matrix of polymer or copolymer fibres (polyethylene terephthalate;PET, polystyrene, polyethylene etc.), acrylic resin and cotton to generate the 3D culture model. The density of the fabric ranges between 10 gm/m^2^ and 50 gm/m^2^. The thickness of the fibers is 0.05-5mm[11].

### 2D cell culture

For 2D culture, cells were seeded at around 60-80% confluency i.e. approximately 0.8 × 10^6^ in 60 mm dish. After cells became 100% confluent, media was removed and cells were washed with PBS, trypsinized and centrifuged at 1000 rpm for 5 minutes. Finally, cells were resuspended in an appropriate volume of respective media and plated in 96 well plate (5X10^3^), depending upon the experiment.

### 3D cell culture

Tissueoids were formed on AXTEX-4D™ platform by using hanging drop methods. Briefly, cells were seeded at around 60-80% confluency i.e. approximately 0.8 × 10^6^ in 60 mm dish. The cells were washed with PBS at 100% confluency, trypsinized and centrifuged at 1000 rpm for 5 minutes. Finally, the process of hanging drop formation was initiated only when the viability of the cells was greater than ≥90%. Cell suspension was made in such a way that 1 ml of respective media contain 2.5 × 10^5^ cells so that 20μl of the media contained a cell number of 5000 cells per drop. The drop was pipetted onto the inner surface of a lid filled with PBS at the bottom. After 24-48 hrs, the inner lid was inverted, and the drops were poured on AXTEX-4D™ platform for tissueoids formation in desired format and analyzed for further experiments attachment and growth (supplementary figure 1).

### T-cell Migration, infiltration and confocal microscopy

T-cells were isolated from human PBMCs and stimulated with CD3 and CD28 for 4 days at 37°C in a humified chamber containing 8% CO2. Before seeding for the migration assay, blood T-cells (0.5 × 10^5^ cells) were added to each insert of a Corning® HTS Transwell®-96 Tissue Culture System (Corning Cat. No. 3387) and allowed to migrate (24 h, 48h and 72h) toward MCF-7 tissueoids. Migration was quantified by cell count using Trypan blue dye. For infiltration, T-cells/Jurkat T-cells were stained by incubation with CFSE green (Thermo Fischer-C34554) in 1:1000 dilution for 1 h. Similarly, MCF-7 tissueoids were stained with Hoechst dye 50 ng/ml for 5 minutes. T-cells/ Jurkat T-cells were allowed to migrate towards 10% FBS and SDF-1α for 48 hrs. Cells were fixed for 5 min in 3.7% formaldehyde solution, washed and mounted with prolong gold antifade. Imaging was performed using confocal microscopy Nikon (A1 R HD 25) at 10X objective. Images were acquired by using Leica TCS SP 8 confocal microscope at 10 X air objective lens, Hoechst was acquired by exciting 405 nm of laser and CFSE was acquired by exciting 488 nm of laser. To avoid inter-channel mixing (405 nm and 488 nm), pictures were captured separately with individual laser and were merged later.

### Cell viability Assay

Cells were cultured in AXTEX-4D™(Tissueoid of 5000 cells/well, n=3) platform in 96 well format. Next, they were treated with 25μg/ml of Nivolumab (PD-1 inhibitor: NP-25) and 2.5μg/ml of Atezolizumab (PD-L1 inhibitor: AL-2.5) along with stimulated as well as unstimulated T cells for 48 hours. Finally, Prestoblue assay has been done as described elsewhere[34]. PrestoBlue (Thermo Fischer; A13261) is a resazurin-based solution for rapidly quantifying the metabolic active cells, providing a metric of cell viability. Briefly 1/10th volume of prestoblue reagent was directly added to the control and treated cells and incubated for 2 hours at 37 °C. Similarly, cell viability was also measured for stimulated and unstimulated T-cells at three time periods (24,48 and 72 hrs) by Prestoblue resazurin based solution. The metabolic rates were measured by the amount of relative fluorescence unit (RFU) at Excitation 560 and Emission 590nm using a spectrofluorometer plate reader (SPECTRA MAX GEMINI EM, Molecular Devices)

### Flow cytometry

Flow cytometric analysis was used to determine the cell surface expression of PD-1 on T cells and PD-L1 on the A-375 cell lines following Nivolumab and atezolizumab treatment. Cells were trypsinized with 0.25% trypsin (Life Technologies) and collected in a complete medium. Cells were washed in phosphate-buffered saline (PBS), re-suspended in flow cytometry staining buffer (FACS) buffer containing 1:100 dilution of target-specific antibodies (anti PD1:cat≠329905; Biolegend, and anti-PDL1: Cat≠329705; Biolegend) and incubated for 30 minutes at 4°C in the dark. After centrifugation and two times washing with FACS buffer, data were acquired using ACEA Novo flow cytometer

### Cytokine profiling

Supernatants were collected from Immune T-cells at various time points (24, 48 and 72hrs) and from NP-25/AL-2.5 treated 3D tumor tissueoids or AL-2.5 treated 2D A375 cells. Cytokine staining for IFN-γ was performed as per the manufacturer instructions (Human IFN-γ ELISA set (RUO), BD Biosciences, USA Cat No. 555142).

### Detection of Cell cytotoxicity using MTT

Cells were cultured in 2D (5000/well, n=3) and AXTEX-4D™(Tissueoid of 5000 cells/well, n=3) platform in 96 well format. Next, they were treated with 2.5μg/ml of Atezolizumab (PDL1 inhibitor: AL-2.5) along with stimulated T cells for 48 hours. Further 20μl MTT solutions from the Stock (5 mg/ml) was added and cells were incubated in CO2 incubator in the dark for 2 hrs. The medium was removed, and formazan crystals formed by the cells were dissolved using 100 μl of DMSO followed by transfer in 96 well plate. The absorbance was read at 570 nm using 630 nm as reference wavelength on a Multiwell plate reader (Biotech Instruments, USA).

## Author Contributions

AB: Planning and execution of experiments, analysis, and writing. SS: Execution and data acquisition and data analysis. BPDP and SK: Planning and execution of experiments. SM: Concept design and reviewing the manuscript. RG analyzed the data with input from all authors and drafting of the manuscript. NMA and PKK: Study concept, design, interpretation of data, providing scientific inputs, obtained funding, and reviewing of the manuscript.

## Acknowledgements

The authors thank Mr. Manish Kumar, IGIB Delhi, and Mr. Ashish Pandey, RCB, Faridabad for confocal microscopy.

---

**Supplementary Figure 1.**
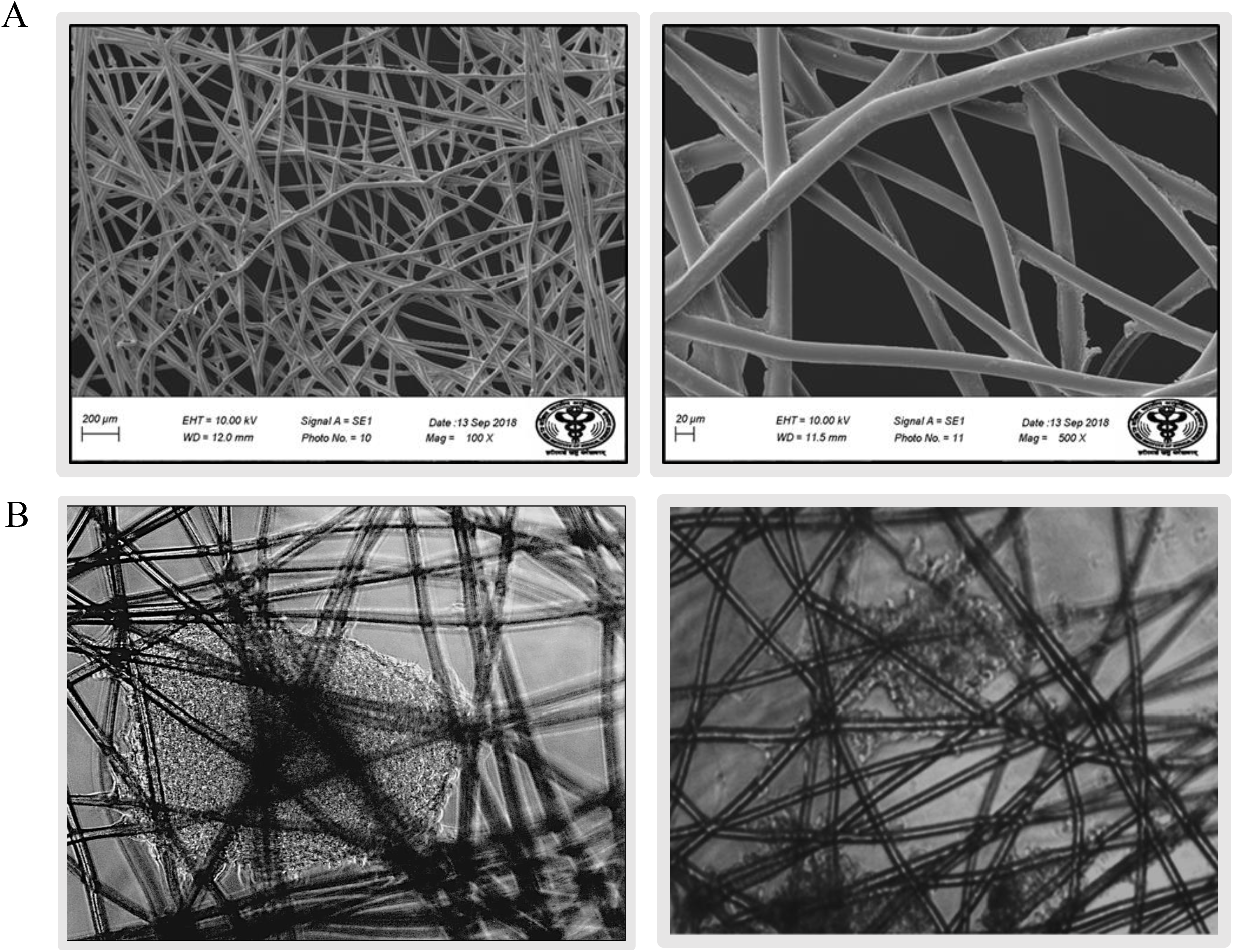
A) Scanning electron microscopy images of AXTEX 4D at 100X (left panel) magnification and provides a larger magnification (500X) (right panel) B) Phase contrast microscopy images of breast (MCF-7:Left panel) and skin (A-375: right panel) cancer 3Dtumor tissueoids grown in 96-well plates onAXTEX-4D™ platform. scale bar = 200μm

